# Cyclodextrin triggers MCOLN1-dependent endo-lysosome secretion in Niemann-Pick type C cells

**DOI:** 10.1101/401190

**Authors:** Fabrizio Vacca, Stefania Vossio, Vincent Mercier, Dimitri Moreau, Shem Johnson, Jonathan Paz Montoya, Marc Moniatte, Jean Gruenberg

## Abstract

In specialized cell types, lysosome-related organelles support regulated secretory pathways, while in non-specialized cells, lysosomes can undergo fusion with the plasma membrane in response to a transient rise in cytosolic calcium. Recent evidence also indicates that lysosome secretion can be controlled transcriptionally and promote clearance in lysosome storage diseases. In addition, evidence is also accumulating that low concentrations of cyclodextrins reduce the cholesterol storage phenotype in cells and animals with the cholesterol storage disease Niemann-Pick type C, via an unknown mechanism. Here, we report that cyclodextrin triggers the secretion of the endo/lysosomal content in non-specialized cells, and that this mechanism is responsible for the decreased cholesterol overload in Niemann-Pick type C cells. We also find that that the secretion of the endo/lysosome content occurs via a mechanism dependent on the endosomal calcium channel MCOLN1, as well as FYCO1, the AP1 adaptor and its partner Gadkin. We conclude that endolysosomes in non-specialized cells can acquire secretory functions elicited by cyclodextrin, and that this pathway is responsible for the decrease in cholesterol storage in Niemann-Pick C cells.

## INTRODUCTION

Niemann-Pick type C (NPC) is an autosomal recessive lysosomal storage disorder (LSD) characterized at the cellular level by an accumulation of cholesterol and other lipids in late endocytic compartments (Vanier, 2010). In NPC, the traffic of LDL-derived cholesterol from lysosomes to the endoplasmic reticulum is defective, which impairs not only cholesterol esterification, but also the feedback transcriptional regulation of cholesterol metabolism via the sterol regulatory element-binding protein (SREBP) pathway (Kristiana et al., 2008; Liscum and Faust, 1987; Pentchev et al., 1985). The disease is caused by loss-of-function mutations in either one of the two late endosome/lysosome proteins NPC1 (Carstea et al., 1997), a multispanning protein of the limiting membrane, and NPC2 (Naureckiene et al., 2000), a globular protein present in the lumen. While solid evidence show that both proteins bind cholesterol (Infante et al., 2008; Xu et al., 2007), structural and mutagenesis data suggest that cholesterol is transferred from NPC2 to NPC1, thereby facilitating export from the organelle(Gong et al., 2016; Kwon et al., 2009; Li et al., 2016; Wang et al., 2010). However, the precise mechanism of export remains unclear. Similarly, it is not clear how cholesterol is transferred from late endosome/lysosomes to reach the regulatory pool in the ER. Several direct routes have been proposed (Pfisterer et al., 2016), but recent findings indicate that the plasma membrane may be the primary destination of LDL-derived cholesterol before reaching the ER(Infante and Radhakrishnan, 2017). This view is consistent with the earlier notion that cholesterol rapidly equilibrates between the plasma membrane and the regulatory ER pool(Lange et al., 2004).

There is no approved treatment against NPC with the exception of Miglustat, which delays but does not arrest the progression of the disease (Patterson et al., 2015). Over the last decade, cyclodextrins are emerging as a possible therapeutical strategy. These cyclic oligosaccharides clear cholesterol storage and reestablish the feedback regulation in cultured cells and NPC mice (Abi-Mosleh et al., 2009; Liu et al., 2010; Rosenbaum et al., 2010). They also improve neurological symptoms and survival in NPC animal models (Liu et al., 2009; Vite et al., 2015), and were shown to decrease the neurological progression of the disease in phase 1-2 trials in NPC patients (Ory et al., 2017). However, the mechanism of cyclodextrin action remains poorly understood.

In recent years, it has become apparent that secretory endosomes or lysosomes (Marks et al., 2013) may play an important role in LSDs. Indeed, activated or overexpressed transcription factors of the TFEB-family can revert storage in Pombe disease and other LSDs by stimulating the secretion of the endo/lysosome content (Martina et al., 2014; Medina et al., 2011; Samie and Xu, 2014). This pathway is distinct from the acute and transient calcium-induced process involved in plasma membrane repair (Andrews et al., 2014; Laulagnier et al., 2011; Reddy et al., 2001). Similarly, storage in cells from NPC and other LSDs can be reverted by δ Tocopherol treatment (Xu et al., 2012) or activation of BK channels (Zhong et al., 2016), which also promote endo/lysosome-mediated secretion. Interestingly, the secretion of endo/lysosome storage materials seems to depend on the activation of the lysosomal cation channel mucolipin-1 (MCOLN1), which is itself responsible for the LSD mucolipidosis type 4 when mutated (Boudewyn and Walkley, 2018). Endosome-or lysosomes-mediated secretion is impaired in MCOLN1 mutant cells (LaPlante et al., 2006), while activating mutations in the channel increase the rate of this exocytic pathway (Dong et al., 2009).

Here, we provide direct evidence that hydroxypropyl-cyclodextrin (HPCD) promotes the secretion of the endo/lysosome content via a mechanism that requires MCOLN1. Our data indicate that the process also depends on FYCO1 — a protein involved in endo-lysosome motility towards the periphery (Pankiv et al., 2010; Raiborg et al., 2015) — as well as the AP1 adaptor and its partner Gadkin, which have both been implicated in the secretion of endo-lysosomes (Laulagnier et al., 2011). Finally, we show that this pathway mediates the cyclodextrin-induced decrease in cholesterol storage within late endocytic compartments of NPC cells.

## RESULTS

### Cyclodextrin reverts the NPC phenotype

NPC1 was depleted with siRNAs in HeLa cells, to provide an experimental system readily amenable to biochemical and cellular manipulations and with a reduced danger of adaptive drift that may occur in patient cells. NPC1 KD cells exhibited the characteristic hallmarks of NPC. Indeed, cholesterol accumulated in late endosomes as revealed by fluorescence microscopy (Fig 1A) and subcellular fractionation (Fig 1D), resulting in increased levels of total cellular cholesterol (Fig 1D). As expected, KD cells also exhibited a reduced capacity to esterify LDL-derived cholesterol (Fig 1B) and to regulate the transcription of cholesterol-dependent genes like the LDL receptor (Fig 1C). Treatment with a low (0.1%) dose of 2-hydroxypropyl-β-cyclodextrin (HPCD) was able to correct the effects of NPC1 KD on cholesterol subcellular distribution (Fig 1A and D), esterification (Fig 1B) and transcriptional regulation (Fig 1C), while having essentially no effect in mock-treated control cells (Fig 1). While HPCD reduced endosomal cholesterol in NPC1 KD cells (Fig 1A and D), the drug only marginally, if at all, affected total cellular cholesterol. Altogether, these observations strengthen the notion that HPCD is able to revert the NPC phenotype in KD cells, presumably by redistributing cholesterol from endosomes to other membranes (Abi-Mosleh et al., 2009; Rosenbaum et al., 2010).

**Figure 1.**
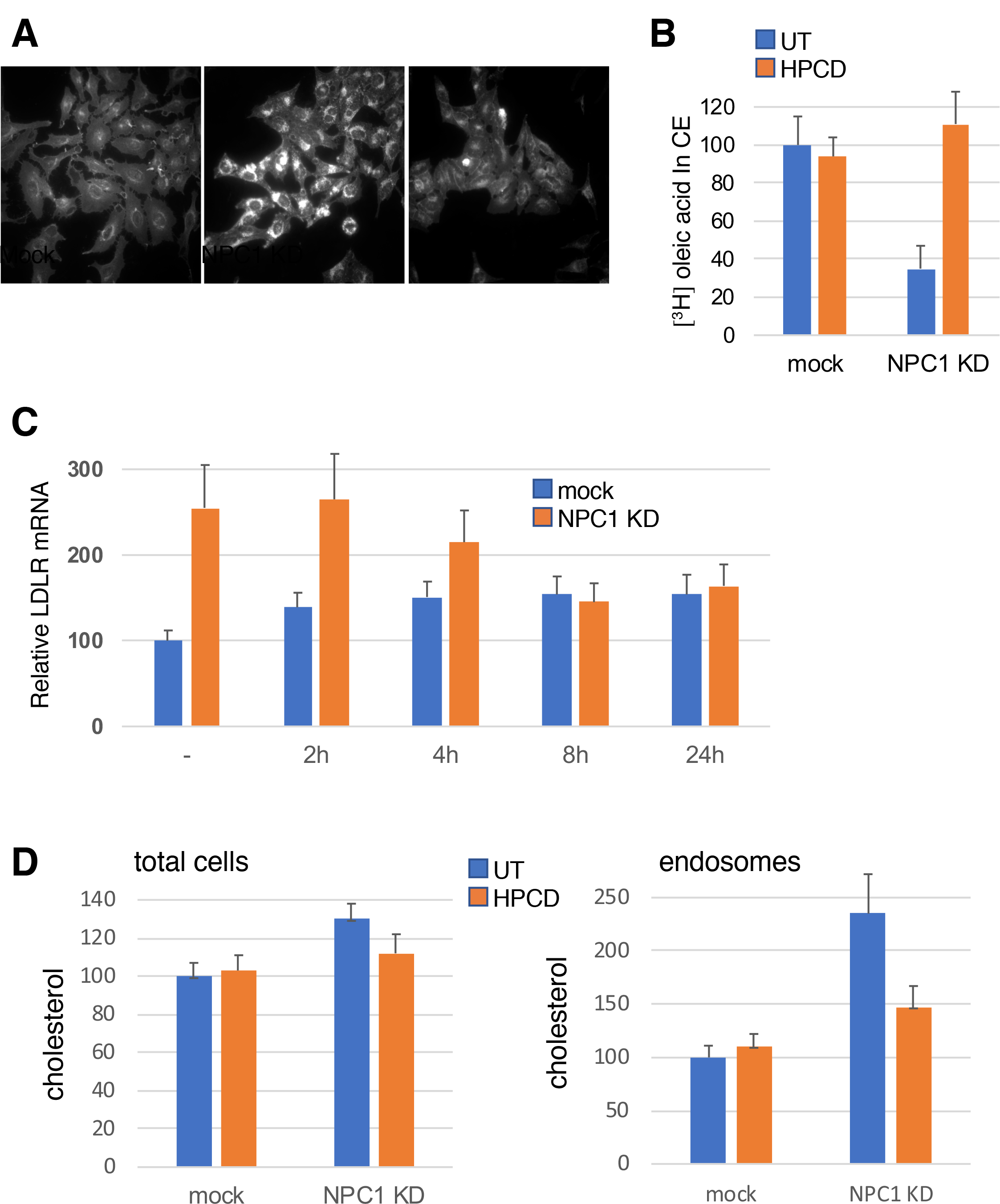
**(A-C)** Hela cells were mock-transfected or transfected with anti-NPC1 siRNA for 72h. When indicated in A, cells were treated for the last 24h with 0,1% HPCD, fixed and stained with filipin. In B, 36h after transfection, cells were starved for 24h in 10% LPDS, then labelled with [^3^H] OA for 12h in 10% FCS. When indicated HPCD 0,1% was added 12h after starvation and maintained during OA incorporation. In C, cells were treated before the end of the 72h with HPCD 0,1%, as indicated. Total mRNA was extracted and LDLR mRNA quantified by RT-PCR. (**D)** BHK cells were transfected as in (A) and treated or not for the last 24h with HPCD 0,1%. A late endosomal fraction was prepared and unesterified cholesterol was quantified (total cells and fractions) enzymatically. B-D: data expressed as a percentage of the mock control.

### HPCD stimulates endo-lysosome secretion

We previously found that the intralumenal vesicles of multivesicular late endosomes contain lysobisphosphatidic acid (LBPA)/ bis(monoacylglycero)phosphate (BMP), an atypical phospholipid that is not detected in other organelles (Kobayashi et al., 2002; Kobayashi et al., 1998). LBPA is functionally linked to cholesterol, since interfering with LBPA phenocopies NPC (Chevallier et al., 2008; Kobayashi et al., 1999). Strikingly, intracellular levels of LBPA were strongly decreased in HPCD-treated cells (40 % of control after 18h), while other phospholipids remained essentially unaffected (Fig 2A). Cellular phospholipids released in the medium could be unambiguously quantified after metabolic labeling with ^32^P. Concomitant with decreased cellular levels, ^32^P-LBPA was released into the culture medium, reaching 2.5-3 % of the total cellular content within 1h, when compared to <1% for other ^32^P-phospholipids (Fig 2B). The analysis ofLBPA released into the medium was not feasible after longer incubations, likely because of its high instability (36).

**Figure 2.**
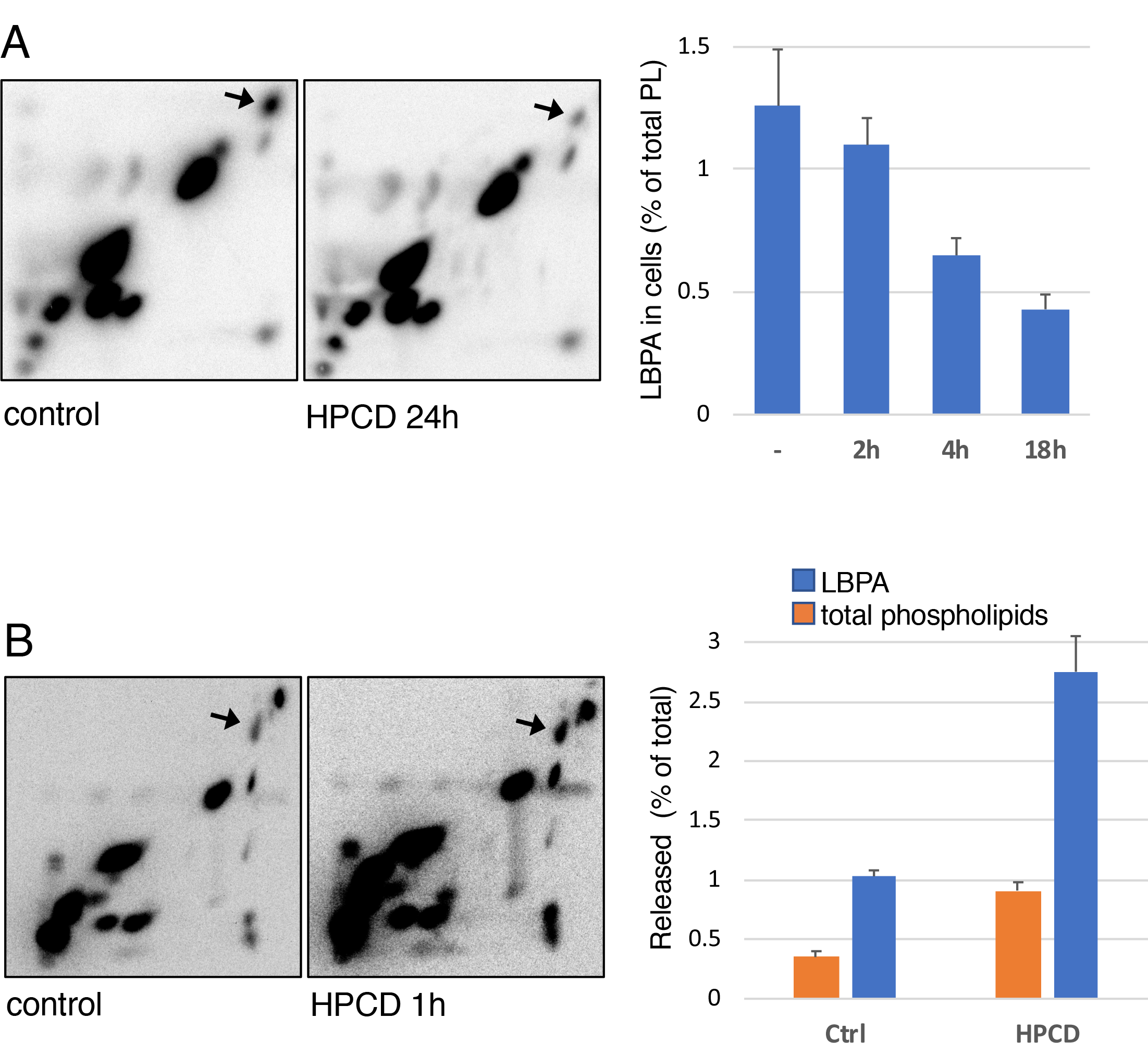
**(A)** Hela cells were labelled with [^32^P] P_i_ for 24h, chased for 14h, then treated or not with HPCD (as in Fig1). Lipid were extracted, separated by 2D-TLC and visualized by autoradiography. The position of LBPA (arrow) was identified by co-migration with pure synthetic LBPA as standard. Lower panel: LBPA quantified in 2D-TLCs at different time after HPCD treatment, and expressed as a percentage of total 32P-phospholipids. **(B)** Cells labelled as in (A) were treated or not with HPCD 0,1% for 1h. The lipids were extracted from medium and separated by 2D-TLC. The lower panel shows the quantification of LBPA and total phospholipids in medium. Data are expressed as a percentage of total cellular LBPA or phospholipids, extracted and quantified in the same experiments

Much like LBPA, treatment with HPCD for 18h strongly reduced the total cellular amounts of the aspartyl endopeptidase cathepsin D (Fig 3A), to ≈60% of controls (Fig 3B). Interestingly, cathepsin D was reduced by HPCD to a similar extent in control (mock-treated) cells and in NPC1 KD cells (Fig 3A-B), indicating that the mechanism controlling cathepsin D release is not affected by NPC1 depletion. In control cells, HPCD also reduced the levels of two other lysosomal hydrolases, β-hexosaminidase and acid lipase (LIPA) to the same extent (Fig 3C) as cathepsin D (Fig 3B). By contrast, the drug had no effect on the total cellular amounts of two endo-lysosomal membrane proteins LAMP1 and NPC1 itself, or on the early endosome marker EEA1 (Fig 3C). Concomitant with decreased cellular amounts, β-hexosaminidase appeared in the medium of HPCD-treated cells in a time-dependent fashion (Fig 4A). This was not due to some deleterious effects of the treatment, since HPCD did not increase the release of the cytosolic enzyme lactate dehydrogenase (LDH). Lysosomal enzyme release occurred with relatively slow kinetics and low efficiency, when compared to acute secretion triggered by a transient raise in cytosolic calcium (Fig 4A). However, in marked contrast to calcium-induced secretion (Laulagnier et al., 2011), the process continued over hours, eventually depleting half of the total cellular amounts after 24h (Fig 3C).

**Figure 3.**
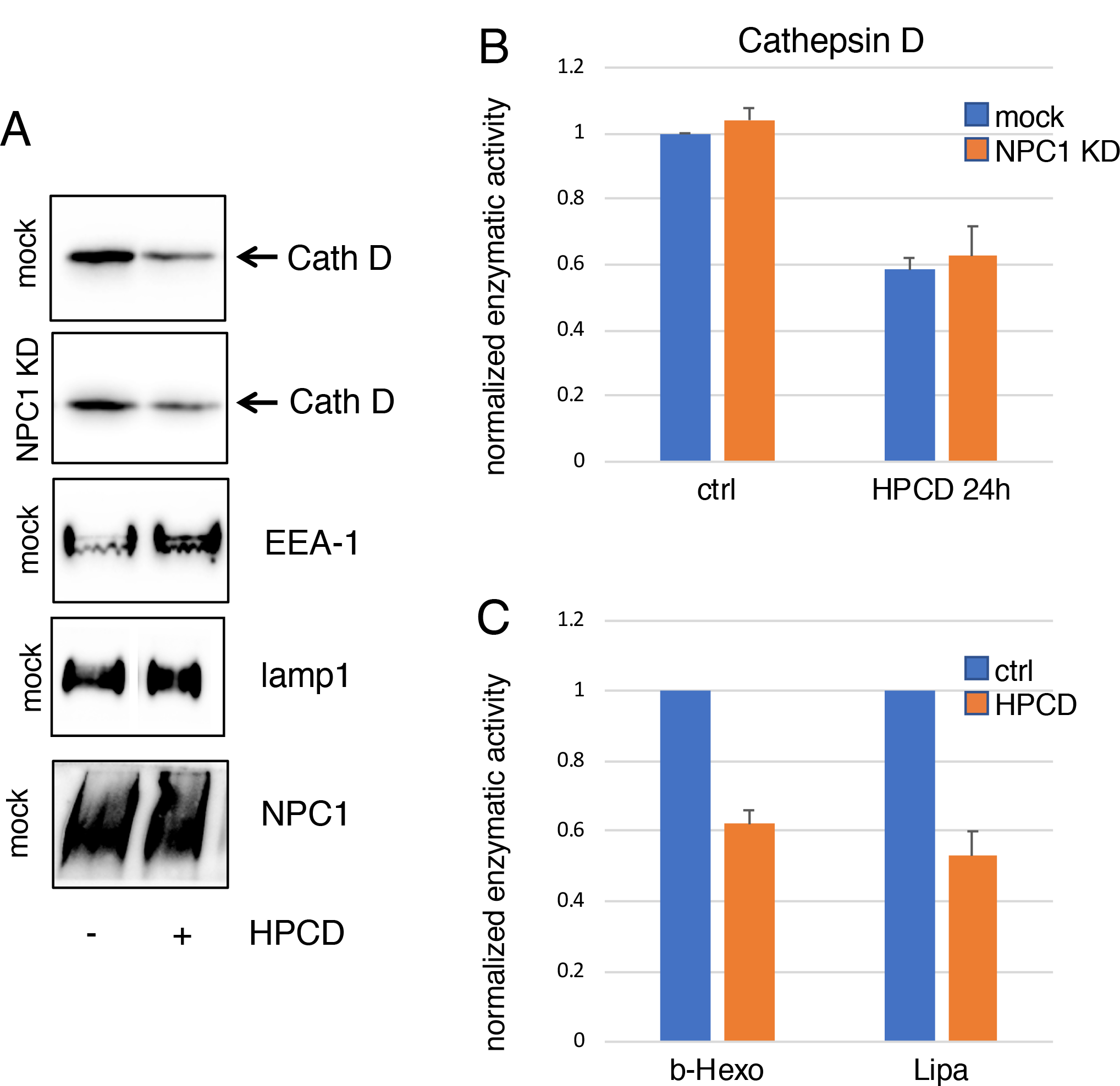
**(A-B)** Hela cells mock-transfected or transfected with anti-NPC1 siRNA for 72h and treated with HPCD (Fig1), and the indicated proteins were analyzed by western blotting in A. Cathepsin D in (A) was quantified (B) by scanning the blots in 3 independent experiments. **(C)** The cellular amounts of acidic lipase (LIPA) and β-Hexosaminidase were quantified enzymatically. Data in (B) and (C) are normalized to the amounts present in the untreated controls.

**Figure 4.**
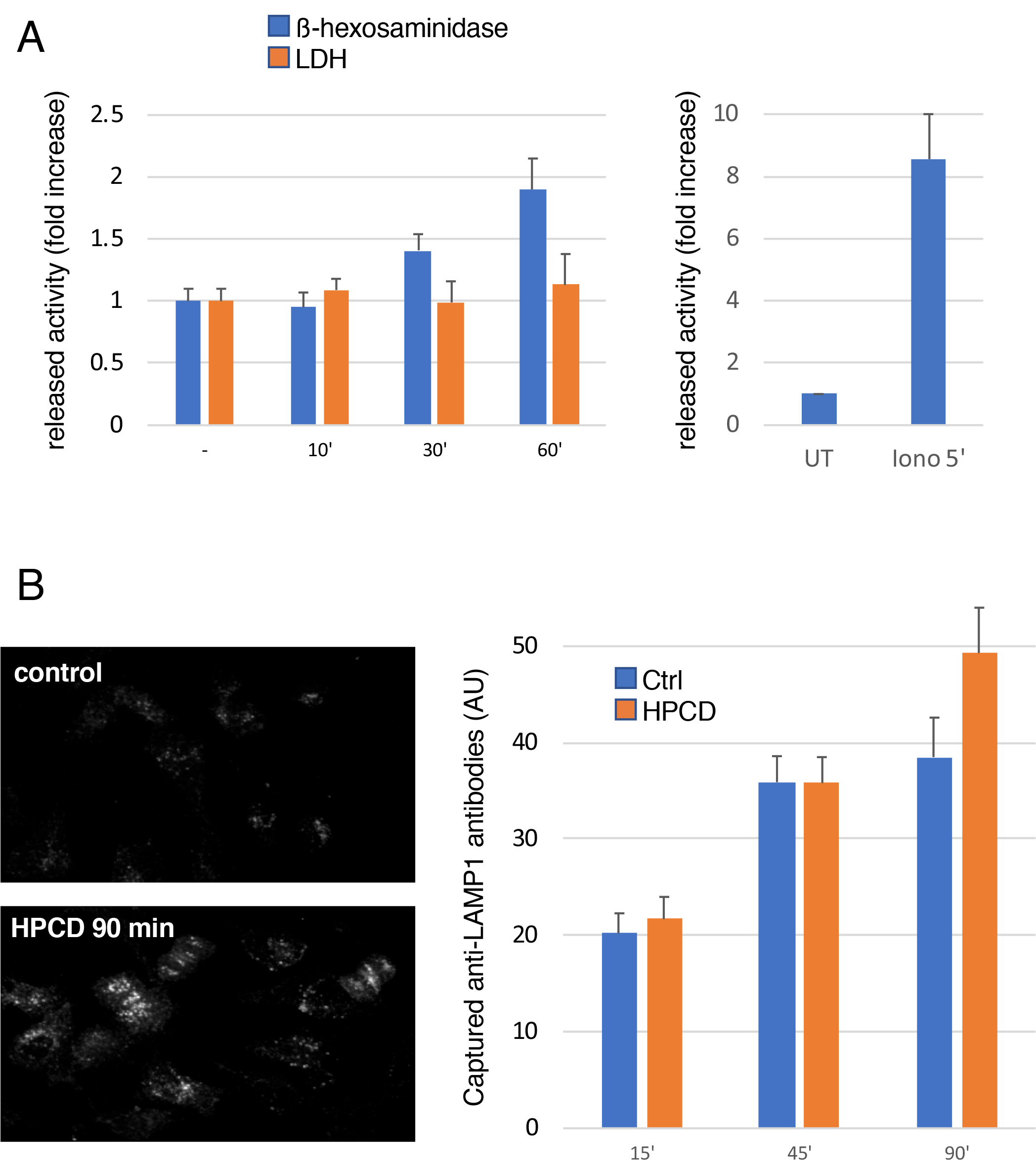
**(A)** Hela cells washed with PBS and then incubated in serum-free medium, were stimulated or not for the indicated time with HPCD 0,1% (right panel) or with ionomycin 5 μM for 5 min (left panel). Then hexosaminidase or lactate dehydrogenase (LDH) activity was measured in the medium and in total cell lysates. The enzyme activity in the medium is normalized to the total activity in lysates. The released enzyme activity is expressed as fold increase compared to untreated cells **(B)** Hela cells in 96-well plates were incubated for the indicated times with lamp1 antibody at 37 °C in the presence or not of HPCD 0,1 %. Cells were than fixed, labelled with Cy3 secondary antibody and analyzed by automated microscopy. The average intensity of the fluorescence signal per cell (arbitrary units) is shown in the right panel.

A three-dimensional analysis by focus ion beam scanning electron microscopy (FIB-SEM) showed that treatment with low concentrations of HPCD for 18h did not affect the morphology of the Golgi complex or mitochondria (Fig EV1) consistent with the notion that HPCD has no detrimental effects on the overall cell organization. However, late endosomes labeled with BSA-gold endocytosed for 4h (at the end of the HPCD incubation period), appeared markedly less electron-dense (Fig 5B and D, low and high magnification), when compared to controls (Fig 5A and C, low and high magnification). Densitometric quantification confirmed that the lumen of HPCD-treated endosomes was significantly lighter than controls (Fig 5E). Presumably, the endosomal content had been released into the medium upon HPCD treatment, fully consistent with our analysis of HPCD-mediated lysosome enzyme release (Fig 4A).

**Figure 5.**
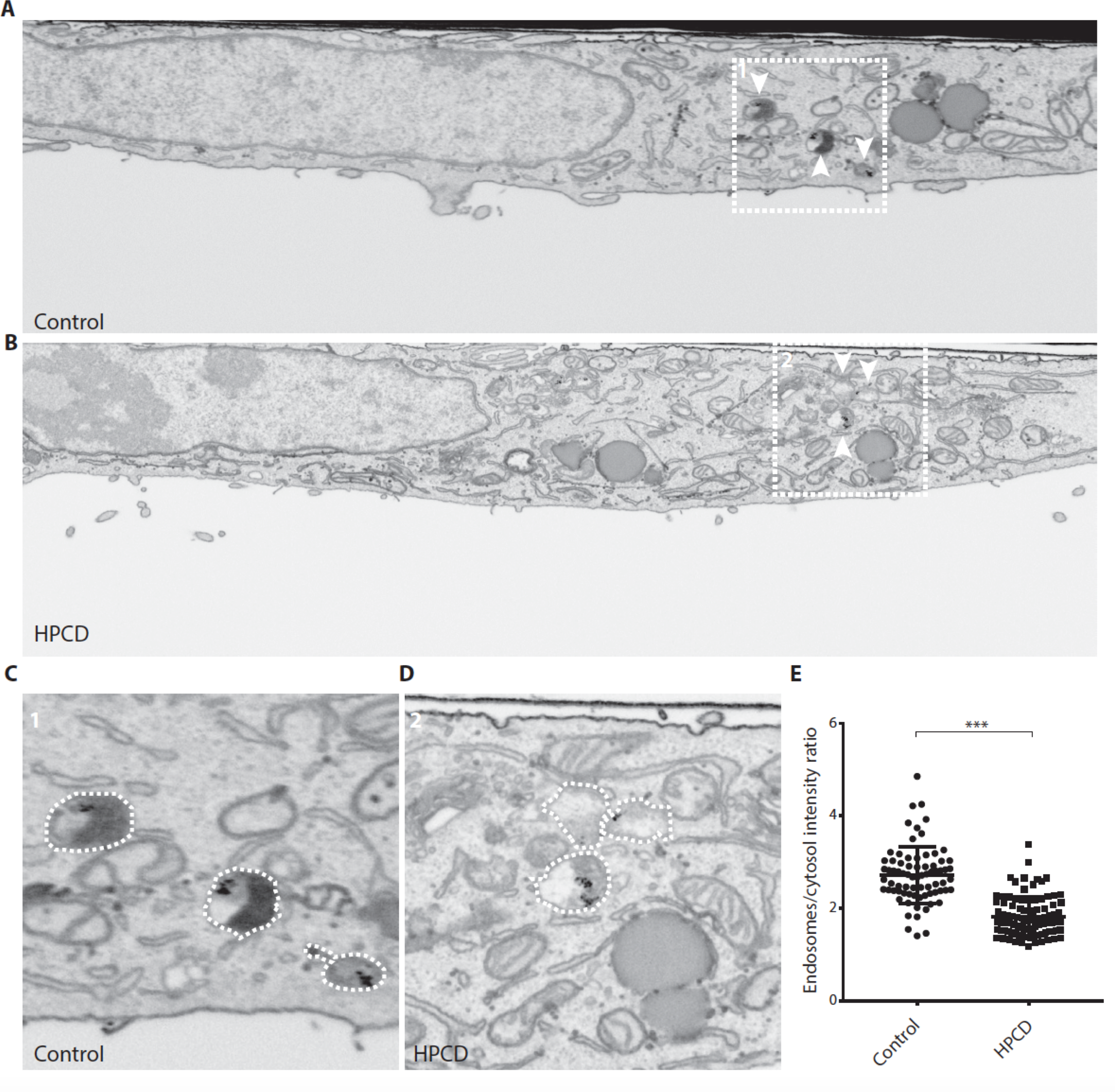
HeLa cells treated (B and D) or not (A and C) with HPCD were incubated with 15nm BSA-gold for 4h at 37°C, and processed for FIB-SEM. The panels show whole-cell slice micrographs (A-B) and high magnification views of the boxed areas (C-D). White arrow point at BSA-gold positive endosomes. The density of endosomes in A-D was quantified by densitometry, and data are expressed relative to the cytoplasm intensity; controls: 2 cells, 74 endosomes; HPCD treated cells: 2 cells, 95 endosomes. ***P<0.0001.

Altogether, these observations suggest that HPCD-induced lysosome enzyme release occurs via endo-lysosome secretion. If so, one may expect the major endo-lysosome membrane protein LAMP1 to appear transiently on the plasma membrane upon endo-lysosome fusion (Laulagnier et al., 2011; Marks et al., 2013). After HPCD treatment, LAMP1 levels hardly changed at the plasma membrane, presumably because LAMP1 was efficiently re-endocytosed (Kornfeld and Mellman, 1989), the secretory process occurring with slow kinetics (Fig 4A). However, in cells incubated with anti-LAMP1 antibodies, HPCD increased the intracellular accumulation of antibodies captured at the plasma membrane after binding to their antigen (Fig 4B). Altogether, these data demonstrate that HPCD triggers the secretion of the endo-lysosome content.

### Mechanism of HPCD-dependent endo-lysosome secretion

In order to better characterize the mechanisms involved in HPCD-mediate endo-lysosome secretion, we used RNAi to deplete proteins involved in related pathways in various cellular systems. These include:

i. RAB27A, secretory pathway of lysosome-related organelles, including pigmentation and immunity (Fukuda, 2013; Marks et al., 2013);
ii. RAB11A, secretion of exosomes (together with RAB27A) derived from intralumenal vesicles of multivesicular endosomes (Savina et al., 2002) (Abrami et al., 2013; Ostrowski et al., 2010);
iii. RAB8A, selective delivery of endosomal cholesterol to the plasma membranes (Kanerva et al., 2013);
iv. RAB3A, lysosome positioning and plasma membrane repair (Encarnacao et al., 2016);
v. syntaxin7 (SYT7) (Reddy et al., 2001), the μ1 subunit of AP1 adaptor and its partner Gadkin (Laulagnier et al., 2011), fusion of endo-lysosomes with the plasma membrane;
vi. the motor protein kinesin-1 (KIF5B) and Arl8A, microtubule-dependent movement of endosomes to the cell periphery (Rosa-Ferreira and Munro, 2011) (Pu et al., 2015);
vii. FYCO1, ER-endosome contact sites and kinesin-mediated motility of lysosomes to the cell periphery (Pankiv et al., 2010; Raiborg et al., 2015);
viii. MCOLN1, an endo-lysosomal calcium channel, exocytosis and clearance of storage materials In LSDs (Martina et al., 2014; Medina et al., 2011).

After depletion by RNAi (Fig EV2), cells were treated with HPCD. Cell-associated cathepsin D was analyzed by western blotting (Fig 6A and Fig EV4) and amounts were quantified (Fig 6B). In cells treated with non-targeting siRNAs, cathepsin D was decreased by HPCD to 50%, as expected (Fig 3). In marked contrast, the amount of cell-associated enzyme was essentially unaffected by HPCD after MCOLN1 depletion, suggesting that lysosome enzyme release was essentially abrogated without this channel. The depletion of the μ1 chain of AP1, gadkin and FYCO1 also reduced cathepsin D secretion but effects were less pronounced, while the depletion of other regulators was without effect. Much like with cathepsin D, HPCD failed to reduce cell-associated LBPA after the depletion of MCOLN1, AP1μ1 or FYCO1, and to a lesser extent after gadkin depletion (Fig 7A). By contrast, HPCD did not affect any other phospholipid, including phosphatidylglycerol (PG), phosphatidylcholine (PC) or phosphatidylethanolamine (PE) whether in control cells or after depletion of any of these factors (Fig 7B). Altogether these data indicate that HPCD-stimulated endo-lysosome secretion is controlled by the calcium channel MCOLN1, together with adaptor protein μ1 and its partner gadkin and by the ER-endosome contact site protein FYCO1 involved in endosome translocation.

**Figure 6.**
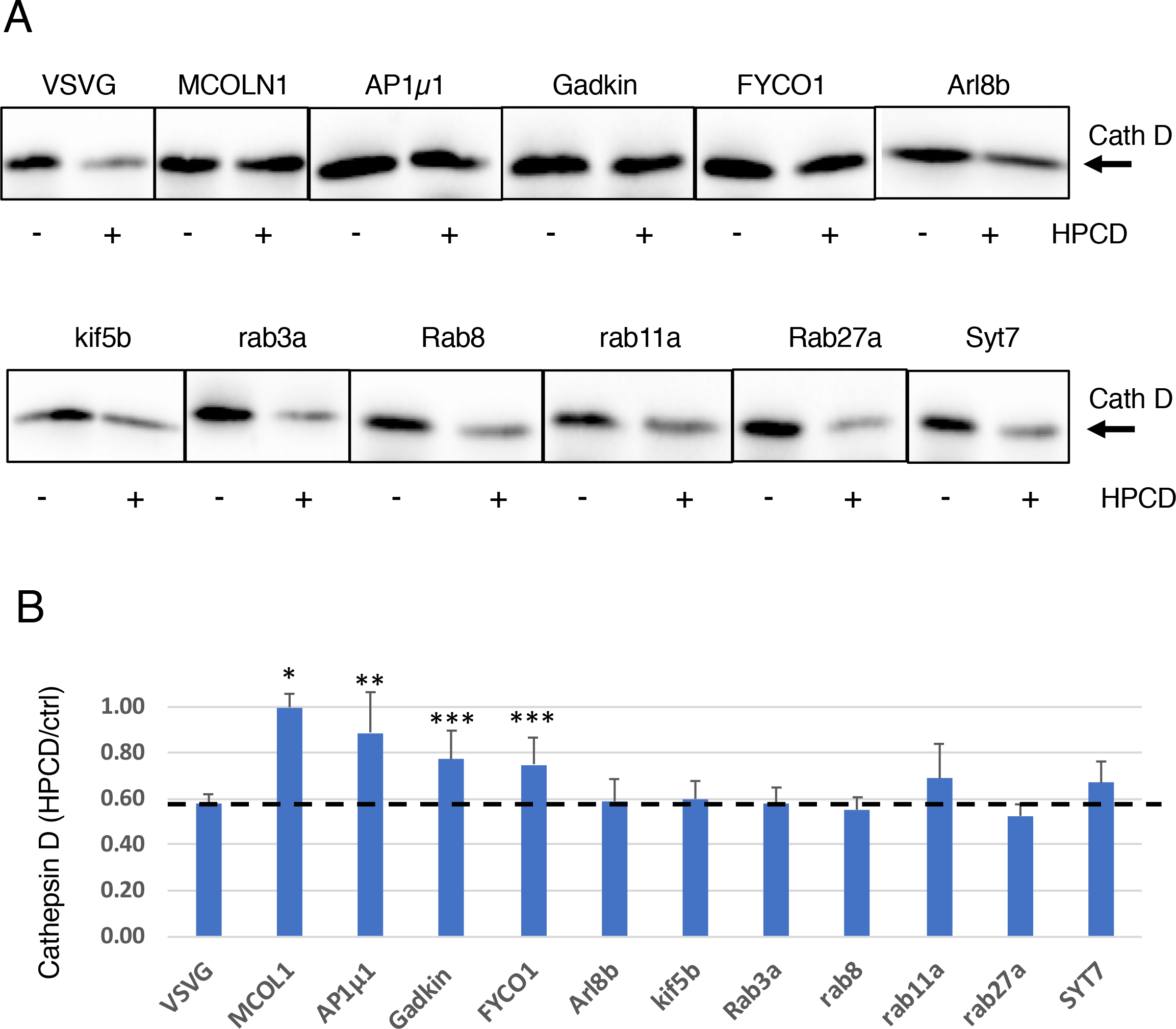
**(A)** Hela cells were transfected with the indicated siRNA for 72h and treated or not with HPCD (as in Fig1). Cells were lysed, and cathepsin D analyzed by western blotting. **(B)** Quantification of cathepsin D normalized to the corresponding β-tubulin signal (shown in Fig. S4). Data are expressed as a ratio of the HPCD-treated samples over the corresponding control for each KD condition (n: 3 independent experiments). *P<0.0001; **P<0.01; ***P<0.02.

**Figure 7.**
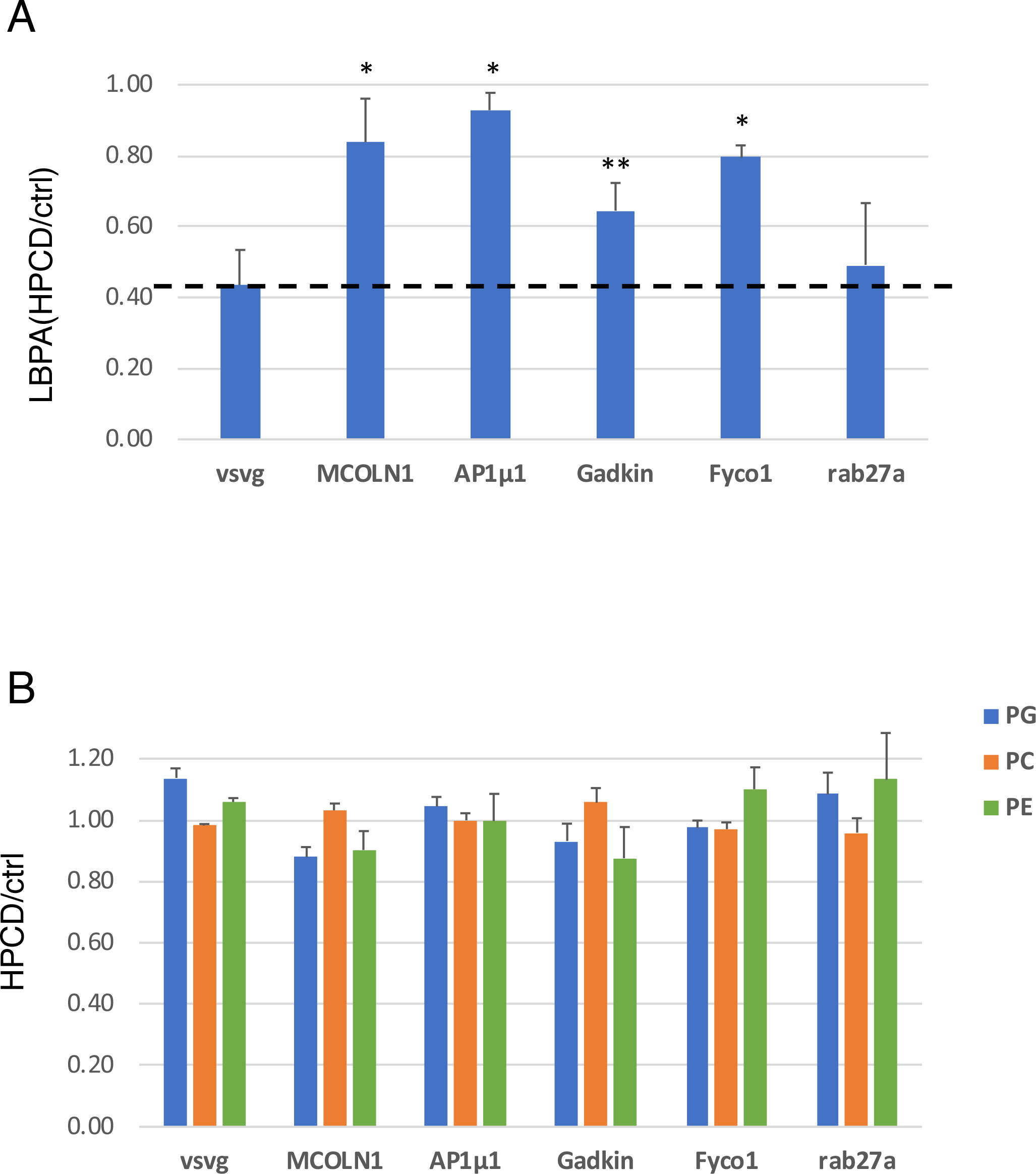
Hela cells were transfected with the indicated siRNA for 72h and treated or not with HPCD (as in Fig.1). The lipids were extracted and quantified by LC-MS. **(A)** All isoforms of LBPA with different acyl chain composition were quantified and normalized to the total phospholipid content. **(B)** All isoforms of phosphatidylglycerol (PG), phosphatidylcholine (PC) and phosphatidylethanolamine (PE) were quantified, normalized as in (A). Data are expressed as aratio of HPCD-treated samples over the corresponding control for each KD condition. (n: 3 independent experiments) *P<0.02; **P<0.05.

### MCOLN1 controls HPCD-dependent endo-lysosome secretion in NPC cells

Next, we investigated whether the same mechanism also mediates the clearance of endosomal cholesterol in NPC KD cells. To this end, we generated a NPC2 knock-out cell line using CRISPR/Cas9, which showed strong cholesterol accumulation in late endocytic compartments (Fig 8A). Treatment of NPC2 KO cells with HPCD massively reduced cholesterol storage (Fig 8A). Quantification by automated microscopy confirmed that, while HPCD did not affect the major endo-lysosomal protein LAMP1 both in NPC2 KO and WT cells, the drug reduced the cholesterol content of LAMP1-containing compartments by half in NPC2 KO cells (Fig 8A and Fig EV3), much like in NPC1 KD cells (Fig 1A).

**Figure 8.**
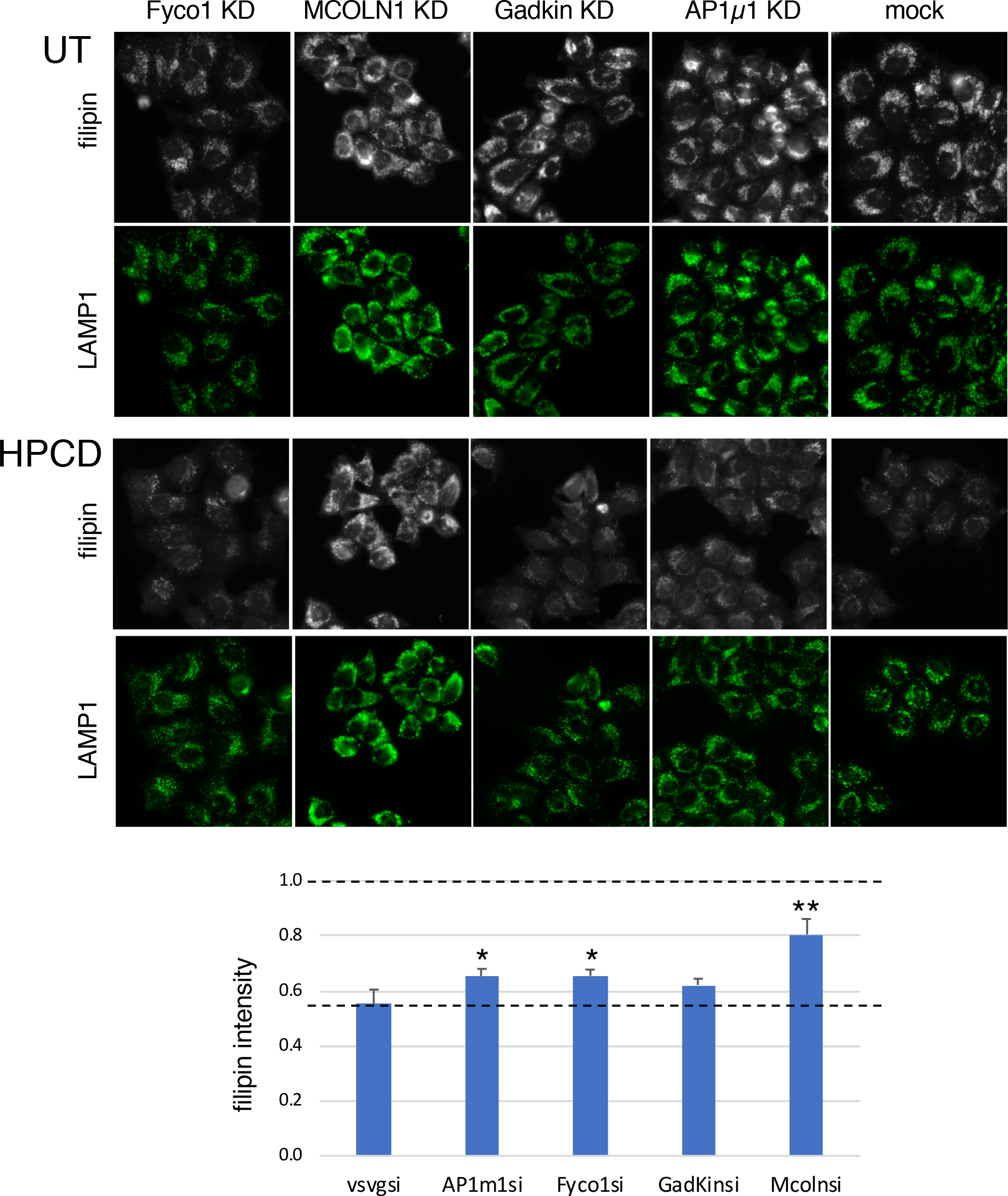
NPC2 KO Hela cells were plated in 96-well plates, transfected with the indicated siRNA for 72h and treated or not with HPCD 0,1% for the last 36h. Cells were fixed, stained with filipin, anti-lamp1 antibodies and propidium iodide (used for segmentation, not shown), and analyzed by automated microscopy. **(A)** Representative images for filipin and lamp1. **(B)** The filipin intensity was quantified in a lamp1 mask and average filipin intensity per cell was calculated. Data are expressed as the ratio between HPCD-treated and untreated cells for each condition. Final averages were calculated over 3 independent experiments *P<0,05; **P<0,005.

The depletion of MCOLN1 by RNAi in NPC2 KO cells had no significant effects on the amounts of LAMP1 or on the cholesterol content of LAMP1-contanining endo-lysosomes (Fig 8A-B and Fig EV3). Remarkably, however, MCOLN1 KD in NPC2 KO cells abrogated the effects of HPCD on cholesterol accumulation (Fig 8A-B and Fig EV3), consistent with the effects of MCOLN1 KD on endo-lysosome secretion in untreated, healthy cells (Fig 6A). Depletion of AP1μ1, gadkin and FYCO1 had a smaller, but significant, effect on HPCD-mediated reduction of cholesterol storage in NPC2 KO cells (Fig 8A, quantification in B). The difference may be due to differential effects of HPCD on cholesterol-laden endosomes in NPC2 cells vs endosomes in control cells. In conclusion our data show that the calcium channel MCOLN1 is directly involved in the regulation of cyclodextrin-mediated endo-lysosome secretion, perhaps together with AP1 and its partner Gadkin, which play a role in calcium-mediated endo-lysosome secretion (Laulagnier et al., 2011) and the ER-endosome contact site protein FYCO1(Pankiv et al., 2010).

## DISCUSSION

In the NPC lysosomal storage disease, mutation of NPC1 or NPC2 causes the accumulation of LDL-derived cholesterol in late endocytic compartments, but the functions of the NPC1 and NPC2 proteins remain incompletely understood (Rosenbaum and Maxfield, 2011). The accumulation of storage materials in NPC endosomes eventually leads to a traffic jam and a collapse of endosomal membrane dynamics (Liscum, 2000; Simons and Gruenberg, 2000), accompanied by defects both in cholesterol movement to the endoplasmic reticulum and transcriptional regulation of cholesterol metabolism (Kristiana et al., 2008; Liscum and Faust, 1987; Pentchev et al., 1985). Compelling evidence now shows that prolonged incubations with low concentrations of cyclodextrins reduces the cholesterol storage phenotype and restores cholesterol-dependent transcriptional regulation (Rosenbaum and Maxfield, 2011) (Vance and Peake, 2011), while cyclodextrin also decreases the progression of neurological symptoms in clinical trials with NPC1 patients (Ory et al., 2017). The mechanism of action of these cyclic oligosaccharides, however, remain poorly understood. Cyclodextrins are thought to act intracellularly after endocytosis as a fluid phase tracer, perhaps bypassing the requirement for the NPC2 protein in the endosome lumen (Rosenbaum et al., 2010). These in vitro studies and our work (this paper) collectively establish that, in the presence of serum, low doses of cyclodextrins do not cause net cholesterol extraction, but instead facilitate redistribution within the cell. This is best illustrated by the restoration of cholesterol esterification and SREBP-mediated feedback regulation in the ER (Abi-Mosleh et al., 2009) (and Fig 1). It has been reported that very short (60min vs. 18-24h in our study) incubations with high doses of cyclodextrin (2% vs. 0.1% in our study) cause calcium-dependent lysosome exocytosis [26]. These high concentrations, however, will cause massive cholesterol depletion from plasma membranes, which may trigger a response to plasma membrane damage (Hissa et al., 2013).

We find that prolonged incubations of control cells with low doses of cyclodextrin causes the secretion of the endo-lysosomal content. Compelling evidence for this secretory pathway comes first from the observed decrease in cellular LBPA content and increase in the medium, since LBPA is only detected in late endosomes (Kobayashi et al., 2002; Kobayashi et al., 1999; Kobayashi et al., 1998). We also find that prolonged incubation with low doses of cyclodextrin causes significant secretion of lysosomal enzymes leading to partial depletion from cells after 18-24 h. Specialized cell types contain lysosome-related organelles (LROs), which have the capacity to undergo fusion with the plasma membrane and to secrete their content in a regulated fashion (Marks et al., 2013) (Luzio et al., 2014). Non-specialized cells also possess the capacity to trigger the acute fusion of endocytic compartments with the plasma membrane in response to a transient rise in cytosolic calcium, presumably as a membrane repair pathway (Laulagnier et al., 2011; Reddy et al., 2001) (Andrews et al., 2014). Given its slow release rate and prolonged duration, our data argue that the cyclodextrin-mediated secretory pathway is distinct from the calcium-mediated pathway and reminiscent of TFEB-induced clearance of lysosomal content (Martina et al., 2014; Medina et al., 2011; Samie and Xu, 2014). This view is also reinforced by our findings that cyclodextrin-mediated secretion is insensitive to the depletion of SYT7 and rab27a, involved in calcium-mediated secretion.

Similarly to TFEB-induced secretion, we find that the secretory pathway elicited by cyclodextrin depends on the endo/lysosomal calcium channel MCOLN1, which is responsible for the LSD mucolipidosis type 4 when mutated (Boudewyn and Walkley, 2018). Previous studies showed that MCOLN1 is involved in the secretion of endo-lysosomal content, since secretion is impaired in MCOLN1 mutant cells (LaPlante et al., 2006), and stimulated by activating MCOLN1 mutations (Dong et al., 2009). It should be noted that is not known whether MCOLN1 function is related to the role of cytosolic calcium in membrane repair. We find that the secretion of endo/lysosomes also depends on the Rab7 effector Fyco1 (Pankiv et al., 2010), involved in endosome translocation to the cell periphery (Raiborg et al., 2015) and in clearance of α-synuclein aggregates (Saridaki et al., 2018). Finally, we find that cyclodextrin-induced secretion of endo/lysosomes also requires the adaptor complex AP1 and its partner protein Gadkin, which are both involved in endo/lysosomes secretion (Laulagnier et al., 2011). Future work will be necessary to unravel the role of MCOLN1 and the mechanism presumably linking this channel to Fyco1 and AP1/Gadkin.

Our data argue that the same mechanism operates in the clearance of storage cholesterol in NPC cells since depletion of MCOLN1 strongly inhibits the HPCD-induced clearance in NPC cells. Depletion of of AP1μ1, gadkin and FYCO1 had a relatively modest, but still significant, effect in NPC cells. One may speculate that the NPC machinery itself plays a role in this secretory pathway, perhaps via ER-endosome contact sites (Raiborg et al., 2015; Rocha et al., 2009). Alternatively, residual lysosomal secretion without these factors suffices to promote cholesterol clearance, perhaps because of some other compensatory mechanisms operating in the NPC KD background. Indeed, MCOLN1 depletion also seems to inhibit more efficiently the cyclodextrin-induced depletion of lysosomal enzymes in WT cells.

In conclusion, our data demonstrate that cyclodextrins trigger the secretion of endo/lysosomes via a pathway dependent on the calcium channel MCOLN1 as well as Fyco1 and the AP1 adaptor and its partner Gadkin. Our data also demonstrate that cyclodextrin clears the endo/lysosomal content of NPC endosomes by stimulating the same MCOLN1-dependent pathway. We conclude that endo-lysosomes in non-specialized cells can acquire secretory functions and that this pathway, elicited by cyclodextrin, is responsible for redistribution of cholesterol and decrease in storage in NPC cells. Our findings also indicate the potential benefit of agents that promote lysosome secretion as possible future strategy to treat Niemann-Pick C and possibly other LSD, given the limitations of currently available therapies.

## METHODS

### Cell Cultures and siRNA transfections

HeLa and BHK-21 cells were maintained as described (Morel and Gruenberg, 2007). HeLa and BHK cells are not on the list of commonly misidentified cell lines maintained by the International Cell Line Authentication Committee. Our HeLa-MZ cells were authenticated by Microsynth (Balgach, Switzerland), which revealed 100% identity to the DNA profile of the cell line HeLa (ATCC: CCL-2) and 100% identity over all 15 autosomal STRs to the Microsynth’s reference DNA profile of HeLa. BHK cells could not be authenticated because no scientific consortium agreed yet on the set of hamster markers and therefore no databank is available to match cell lines. Cells are mycoplasma negative as tested by GATC Biotech (Konstanz, Germany). For siRNA transfection, Hela cells were plated in 6 cm-dishes 6h before transfection in antibiotic-free medium in order to reach 70% confluency. Cells were then transfected with siRNA (140 pmol in 700 μl Optimem) and siRNA max (7 μl in 700 μl Optimem) following the manufacturer’s instructions. 20h after transfection cells were split at a 1:5 dilution. The experiments always ended 72h after transfection.

### Fluorescent automated microscopy

Transfections with siRNAs of HelaMz wt and D11 (crisprNpc2) was with Lipofectamine RNAiMax (Life Technologies AG; Basel, Switzerland) using the manufacturer’s instructions in 96 well-plate from IBIDI (ref 250210) (6000 cells/well). After 36h cells were treated or not with HPDC (0,1 % w/v) for 36h and then fixed in 3% paraformaldehyde for 20 min. Cells were stained with anti-Lamp1 antibody (D2D11-XP rabbit ^®^mAb #9091, Cell Signaling) (1:300) and treated with RNAse (200 μg/ml) followed by propidium iodide (PI) (5μg/ml), filipin (50 μg/ml) and AlexaFluor647 anti-Rabbit secondary antibodies (1:400). Cells were imaged using the ImageXpress Micro XL Confocal automated microscope (Molecular Devices LLC; Sunnyvale, CA). Images were quantified using Meta Xpress images analysis Software. The Custom Module editor software was used to first segment the image and generate relevant masks (propidium iodide for the nucleus, propidium iodide at a lower threshold for the whole cell, LAMP1 for late endosomes/lysosomes), which were then applied on the fluorescent images to extract relevant measurements. The filipin signal was quantified in the lamp1 mask and then expressed as total integrated intensity per cell.

### LAMP1 uptake and analysis

Hela cells were plated in 96 well-plates (10000 cells/well). After 24h, cells were incubated with anti-LAMP1 antibodies (D2D11) (1:300) in full medium with or without HPCD (0,1 %) for 1h, washed 3X with PBS and fixed in 3% paraformaldehyde for 20min. Cells were stained with propidium iodide (5μg/ml) and Cy2-labeled anti-Rabbit secondary antibodies (1:400). Images were acquired with the IXM^™^ microscope (Molecular Devices LLC), and quantified using Meta Xpress Software. Cells were segmented and the total integrated intensity of the LAMP1 signal per cell was measured.

### Cholesterol esterification assay

Cholesterol esterification was measured by incorporation of [^3^H] oleate into newly synthesized cholesteryl oleate as previously described (Chamoun et al., 2013)). Briefly, Hela cells in 6 cm dishes, were starved in 10 % LPDS containing medium for 24h and then labelled for 12h with [9,10–^3^H] OA (45?Ci/mmol, 7.5 μCi/point) complexed with fatty-acid-free BSA, for 14 h in 10% FCS containing medium. After lipid extraction, lipids were separated by thin-layer chromatography (TLC) and [^3^H]-labelled lipids were visualized and quantified using a cyclone Phosphorimager with a tritium Sensitive Screen (PerkinElmer).

### Phospholipid analysis by 2D-TLC

Hela cells were plated in 10 cm dishes, metabolically labelled with [^32^P]P_i_ (80 μCi/point) for 24h and chased for 14h before HPCD addition. At the end of the treatment, PNSs were prepared and lipids extracted with the Folch method (Folch et al., 1957). To analyze the release of phospholipids in the medium, lipids were also extracted from the medium. Lipids were separated by 2D-TLC (Sobo et al., 2007). [^32^P]-labelled phospholipids were visualized and quantified using a cyclone Phosphorimager. LBPA was identified by co-migration with 18:1/18:1 LBPA standard (Echelon Inc, Salt Lake City).

### Cholesterol Quantification

The cholesterol content of cells and sub-cellular fractions was quantified enzymatically using the Amplex Red kit (Molecular probes) (Amundson and Zhou, 1999). Late endosome fractions were prepared from BHK cells by flotation in sucrose gradients (Aniento et al., 1993). Cells or sub-cellular fractions were lysed with lysis buffer (20 mM TRIS-HCl pH 7,5, 150 mM NaCl, 1% Triton X-100) and 5 μg proteins/point (total cells) or 1 μg protein/point were used to quantify cholesterol in a 96-well plate format as described in the kit protocol.

### RT-PCR

Total-RNA was extracted using RNeasy Mini Kit from Qiagen (ref.74104) from monolayers of Hela according to manufacturer’s recommendation. cDNA synthesis was carried out using SuperScript^™^ VILO^™^ cDNA Synthesis Kit (Life Technologies AG; Basel, Switzerland) from 250 ng of total RNA. mRNA expression was evaluated using SsoAdvanced SYBR Green Supermix (Bio-Rad Laboratories, Hercules, CA) with 10 ng of cDNA with specific primers of interest on a CFX Connect real-time PCR Detection System (Bio-Rad). Relative amounts of mRNA were calculated by comparative CT analysis with 18S ribosomal RNA used as internal control. All primers are QuantiTect primer from Qiagen (Hilden, Germany). Primers: ***AP1AR* (QT00054439);** AP1M1 (QT00012215); ARL10C (QT00044758); FYCO1 (QT00037009); KIF5B (QT00046585); MCOLN1 (QT00094234); PLEKHM2 (QT02307109); RAB11A (QT00218491); RAB27A (QT00040054); RAB3A (QT00007490); RAB8A (QT00002485); SYT7 (QT00086975); SYTL4 (QT00088662); RRN18S (QT00199367); LDLR_1 (QT00045864); HMGCR (QT00004081).

### Lipid mass spectrometry

Lipids were extracted as previously described (Scott et al., 2015) with minor modifications. Briefly, cells were grown on 10-cm plastic plates before washing with, and then scraped into, cold PBS followed by centrifugation for 5 min at 300 *g* at 4°C. Cell pellets were resuspended in 100 μl cold water before addition of 360 μl methanol and lipid internal standards (list). Next, 1.2 ml of 2-methoxy-2-methylpropane (MTBE) was added and the samples followed by 1 h shaking at room temperature. A total of 200 μl of water was added to induce phase separation, and the upper phase was collected. Total phosphates were quantified with an ammonium Molybdate colorimetric assay (Loizides-Mangold et al., 2012). Dried lipid samples were re-dissolved in chloroform–methanol (1:1 v/v). Separation was performed using a HILIC Kinetex column (2.6 μm, 2.1 × 50 mm2) on a Shimadzu Prominence UFPLC xr system (Tokyo, Japan): mobile phase A was acetonitrile:methanol 10:1 (v/v) containing 10 mM ammonium formate and 0.5% formic acid; mobile phase B was deionized water containing 10 mM ammonium formate and 0.5% formic acid. The elution of the gradient began with 5%B at a 200μL/min flow. The gradient increased linearly to 50% B over 7min, then the elution continued at 50% B for 1.5 min and finally the column was re-equilibrated for 2.5 min. The sample was injected in 2μL chloroform:methanol 1:2 (v/v). Data were acquired in full scan mode at high resolution on a hybrid Orbitrap Elite (Thermo Fisher Scientific, Bremen, Germany). The system was operated at 240’000 resolution (m/z 400) with an AGC set at 1.0E6 and one microscan set at 10 ms maximum injection time. The heated electro spray source HESI II was operated in positive mode at a temperature of 90°C and a source voltage at 4.0KV. Sheath gas and auxiliary gas were set at 20 and 5 arbitrary units respectively while the transfer capillary temperature was set to 275°C. The mass spectrometry data were acquired with LTQ Tuneplus2.7SP2 and treated with Xcalibur 4.0QF2 (Thermo Fisher Scientific). Lipid identification was carried out with Lipid Data Analyzer II (LDA v. 2.5.2, IGB-TUG Graz University) (Hartler et al., 2011). Peaks were identified by their respective retention time, m/z and intensity. Instruments were calibrated to ensure a mass accuracy lower than 3 ppm. Data visualization was improved with the LCMSexplorer web tool hosted at EPFL (https://gecftools.epfl.ch/lcmsexplorer).

### Analysis by focused ion beam – scanning electron microscopy (FIB-SEM)

HeLa cells treated of not HPCD were incubated with 15nm BSA-gold for 4h at 37°C, rinsed once with PBS, fixed for 3h on ice using 2.5% glutaraldehyde/2% paraformaldehyde in buffer A (0.15M cacodylate, 2mM CaCl_2_). Then cells were extensively washed on ice in buffer A then incubated 1h on ice in 2% osmium tetroxide and 1.5% potassium Ferro cyanide in buffer A and finally rinsed 5 × 3min in distilled water at room temperature. Cells were then incubated 20min at roomtemperature in 0.1M thiocarbohydrazide, which had been passed through a 0.22 μm filter, and extensively washed with water. Samples were incubated overnight at 4°C protected from light in 1% uranyl-acetate, washed in water, further incubated in 20mM Pb aspartame for 30min at 60°C and finally washed in water after this last contrasting step. Samples were dehydrated in a graded series ethanol, embedded in hard Epon and incubated for 60h at 45°C then for 60 hours at 60°C. A small bloc was cut using an electric saw and the bloc was incubated approximatively 30min in 100% xylene in order to remove the plastic left. Finally, the bloc was mounted on a pin, coated with gold and inserted into the chamber the HELIOS 660 Nanolab DualBeam SEM/FIB microscope (FEI Company, Eindhoven, Netherlands). ROI were prepared using focused ion beam (FIB) and ROI set to be approximatively 20μm wide. During acquisition process, the thickness of the FIB slice between each image acquisition was 10 nm. For endosomal density quantification, gold containing endosomes were identified, image intensity inverted and the mean intensity of endosome was measured and divided by the intensity of the cytoplasm for each slice. Results were analyzed using GraphPad Prism. The Gaussian distribution of the data were tested using Kolmogorov–Smirnov test (with Dallal–Wilkinson–Lillie for P value). As it was a non-Gaussian distribution, a two-tailed non-parametric Mann–Whitney U test was used in order to compare conditions.

### β-Hexosaminidase, LDH and Acidic lipase enzymatic assays

β-Hexosaminidase and LDH enzymatic activities were measured from culture supernatant (100μl) total cell lysate (5 μg) or light membrane fractions (2,5 μg) as described (Laulagnier et al., 2011). Acidic lipase activity was measured from light membrane fractions (5 μg) using **4**-Methylumbelliferyl oleate (4-MUO) as substrate (Sheriff et al., 1995). Sample diluted in 25 μl of lysis buffer was mixed with 25 μl substrate solution (4-MUO 2mg/ml in triton X100 4%) and 100μh of assay buffer (NaOAc 200 mM pH 5,5, tween 80 0,02 %). Reaction was incubated for 30-60 min at 37 °C and stopped by adding 100 ml of Tris 1M, pH 8,8. Fluorescence of the reaction product 4-methylumbelliferone (ex. 365 nm, em. 460 nm) was measured in a spectrofluorimeter and compared with a standard curve.

### CRISPR/Cas9 NPC2 KO cell line

Guide sequences to produce NPC2 KO cells were obtained using the CRISPR design tool (Ran et al.,2013):(fwd:TAATACGACTCACTATAGGTCCTTGAACTGCACC;rev: TTCTAGCTCTAAAACAACCGGTGCAGTTCAAGGA). The sequences were used to insert the target sequence into the pX330 vector using Golden Gate Assembly (New England Biolabs) and transfected into cells. Knock-out clones were isolated by serial dilution and confirmed by RT-PCR, Western blotting and filipin staining.

### Other Methods

Protein determination (Bradford, 1976) and gel electrophoresis (Laemmli, 1970) were described. In western blot analysis, primary antibodies were incubated overnight at 4°C, and secondary HRP-linked antibodies for 50min at RT. Antigens were visualized using the westernbright quantum reagent (Advansta, Menlo Park, CA) with the FUSION solo Image Station; band intensities were measured on non-saturated raw images with FUSION solo image analysis software.

## ACKNOWLEDGEMENTS

Support was from the Swiss National Science Foundation, the NCCR in Chemical Biology and LipidX from the Swiss SystemsX.ch Initiative, evaluated by the Swiss National Science Foundation (to J. G.). FV was supported by a fellowship from the National Niemann-Pick C Disease Foundation (NNPDF).

## AUTHOR CONTRIBUTIONS

FV carried out most of the experiments and analyses and wrote the manuscript; SV contributed the mRNA determinations and microscopy; VM carried out the FIP-SEM experiments; DM carried out the experiments by automated microscopy; SM generated the NPC knock-out cell line using CRISPR/Cas9; JPM and MM carried out the analysis of lipids by mass spectrometry; and JG raised the necessary funds, supervised the entire project and wrote the manuscript

## Cyclodextrin triggers MCOLN1-dependent endo-lysosome secretion in Niemann-Pick type C cells

[Fabrizio Vacca, Stefania Vossio, Vincent Mercier, Dimitri Moreau, Shem Johnson, Jonathan Paz Montoya, Marc Moniatte and Jean Gruenberg]

## EXPANDED VIEW

**Figure EV1.**
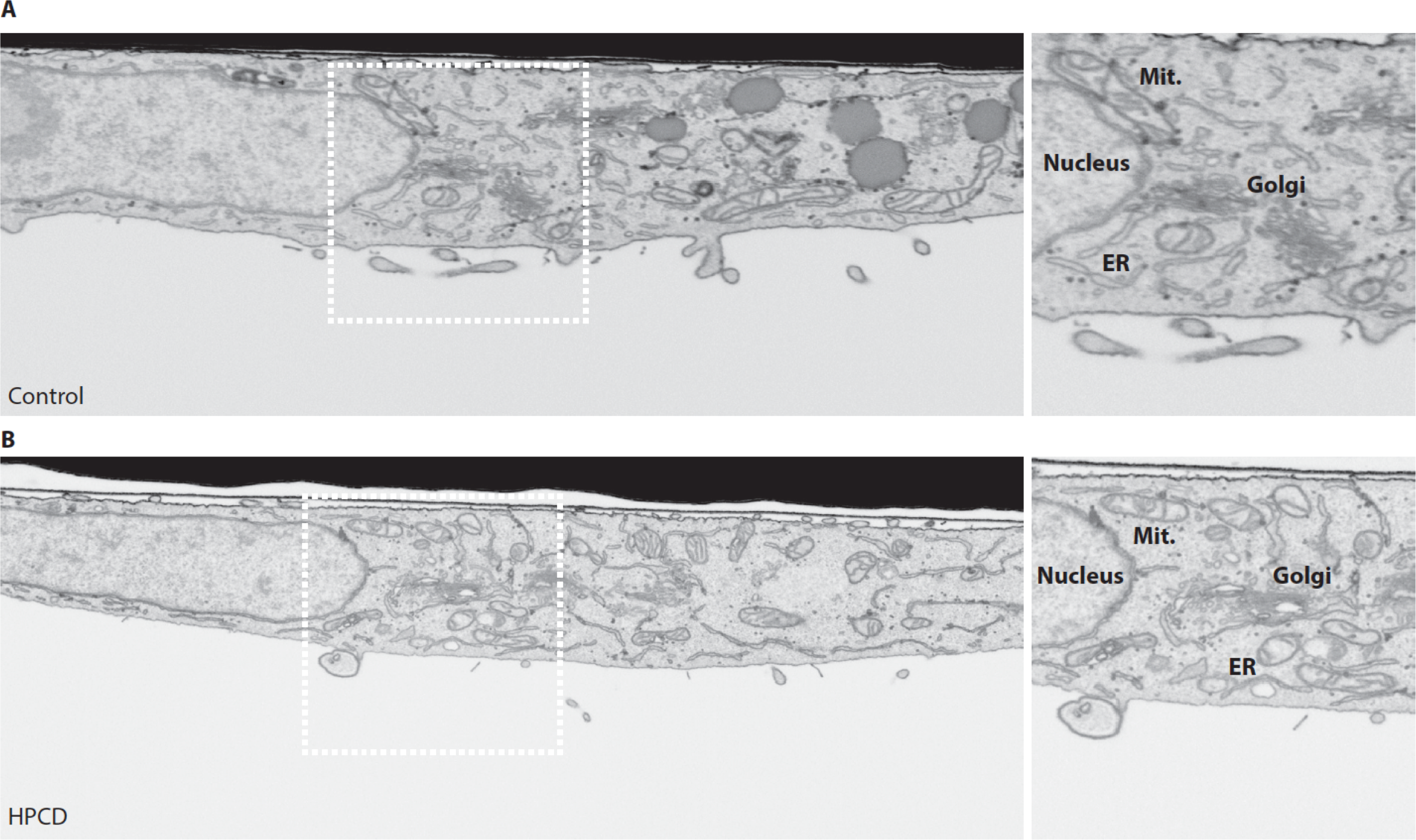
The panels show the same cells as analyzed in Fig 5, which were treated (B) or not (A) with HPCD and processed for FIB-SEM. The panels show whole-cell slice micrographs (A-B) and high magnification views of the boxed areas (C-D) with the nucleus, mitochondria (Mit.), endoplasmic reticulum (ER) and the Golgi apparatus (Golgi).

**Figure EV2.**
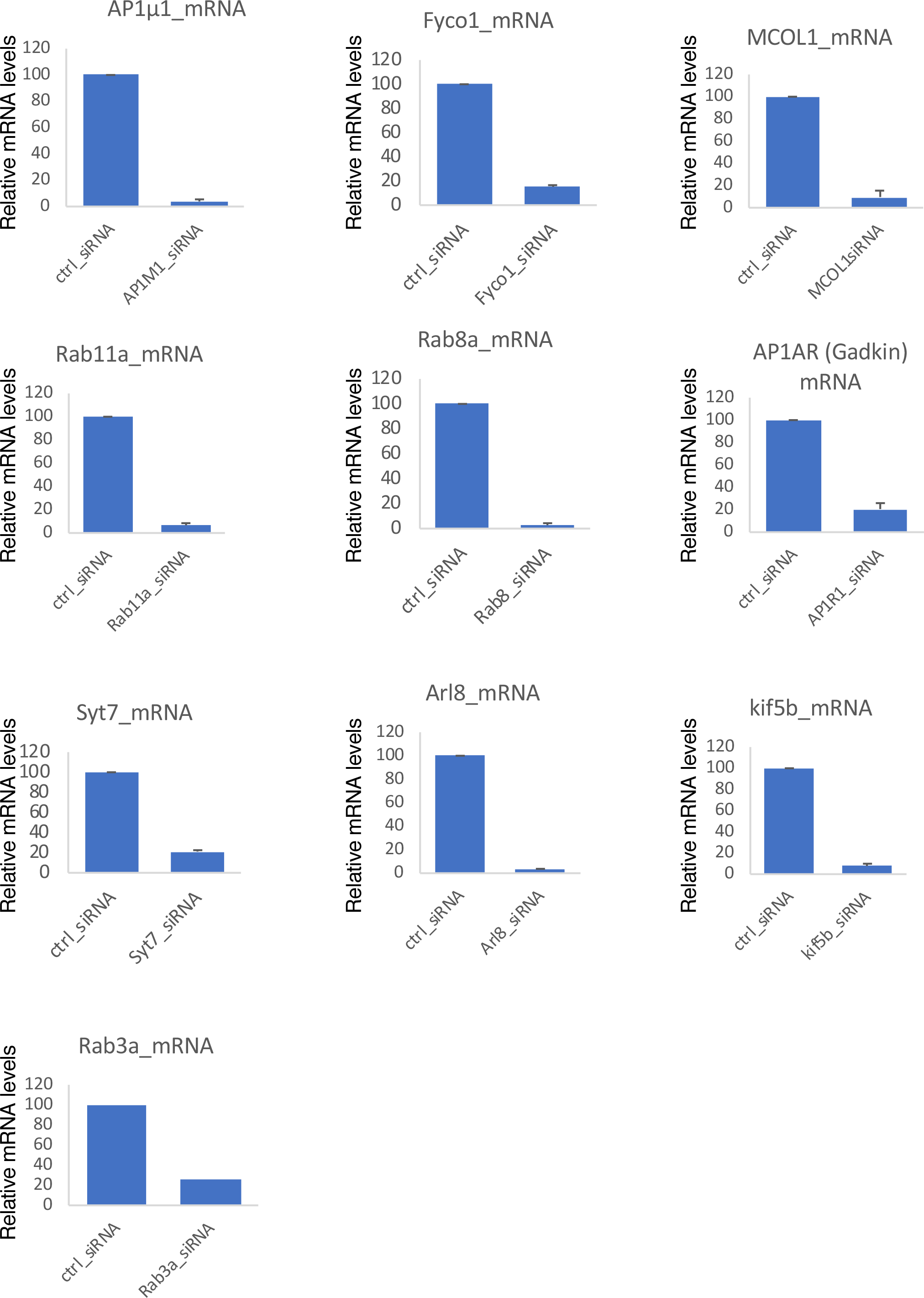
The indicated mRNAs were quantified by RT-PCR with or without depletion by RNAi.

**Figure EV3.**
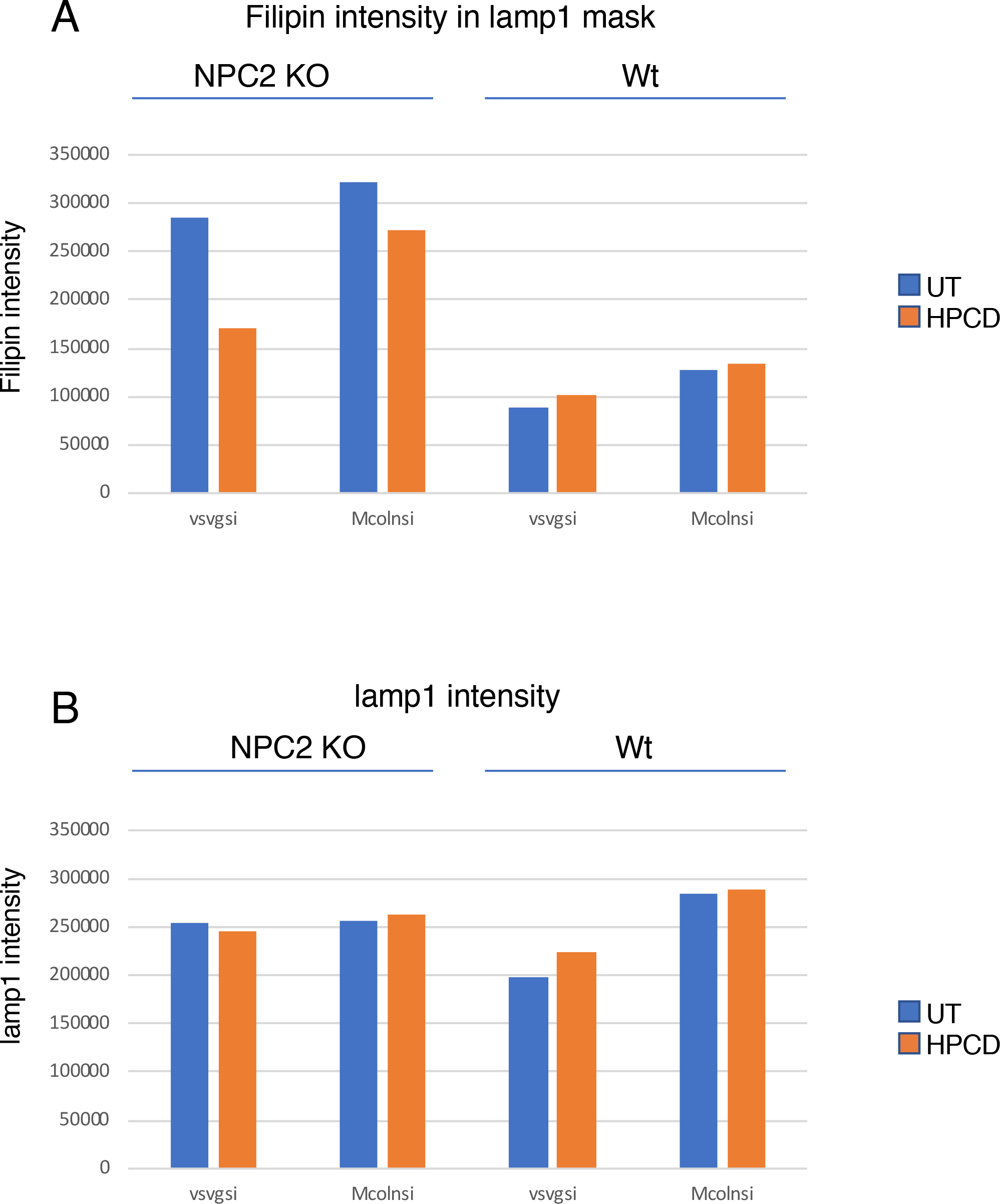
As in Fig 8, NPC2 KO Hela cells were plated in 96-well plates, transfected with the indicated siRNA for 72h and treated or not for the last 36h with HPCD 0,1%. Cells were than fixed and stained with filipin, LAMP1 ab and propidium iodide (used for segmentation, not shown). Images were acquired by automated microscopy. Images were analyzed with Metaxpress software. The filipin (A) and LAMP1 (B) intensity was quantified in the corresponding masks and the average intensity per cell was calculated as in Fig 8 (n= 3 independent experiments).

**Figure EV4.**
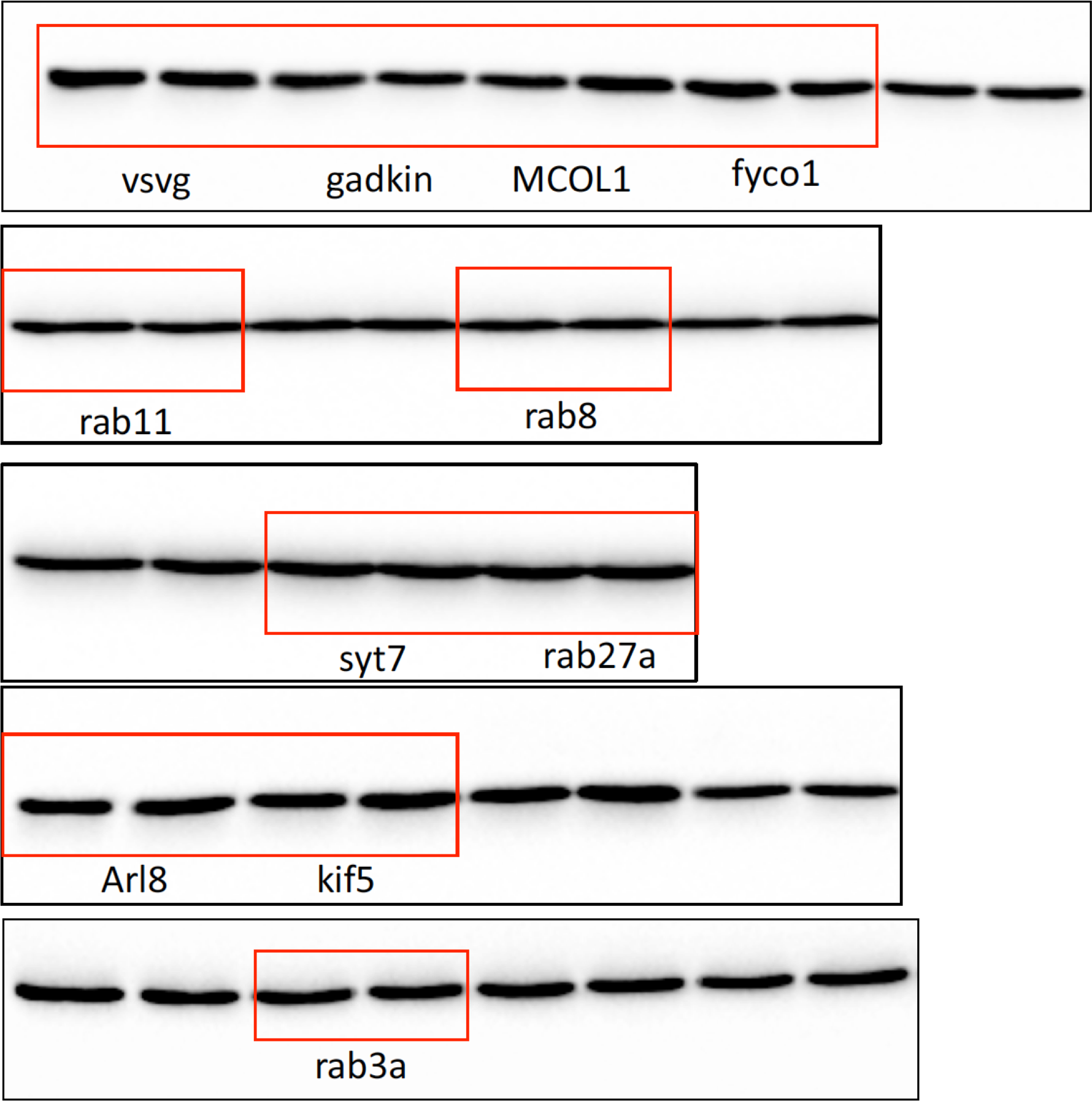
Hela cells were transfected with the indicated siRNA for 72h and treated or not for the last 24h with HPCD 0,1%. Cells were then lysed, and the lysates shown in Fig 6 were analyzed by western blotting using antibodies against tubulin as equal loading marker. The lanes are presented exactly as in Fig 6 for each condition).

## REFERENCES

Abi-Mosleh, L., Infante, R.E., Radhakrishnan, A., Goldstein, J.L., and Brown, M.S. (2009). Cyclodextrin overcomes deficient lysosome-to-endoplasmic reticulum transport of cholesterol in Niemann-Pick type C cells. Proc Natl Acad Sci U S A 106, 19316–19321.

Abrami, L., Brandi, L., Moayeri, M., Brown, M.J., Krantz, B.A., Leppla, S.H., and van der Goot, F.G. (2013). Hijacking multivesicular bodies enables long-term and exosome-mediated long-distance action of anthrax toxin. Cell Rep 5, 986–996.

Amundson, D.M., and Zhou, M. (1999). Fluorometric method for the enzymatic determination of cholesterol. J Biochem Biophys Methods 38, 43–52.

Andrews, N.W., Almeida, P.E., and Corrotte, M. (2014). Damage control: cellular mechanisms of plasma membrane repair. Trends in cell biology 24, 734–742.

Aniento, F., Emans, N., Griffiths, G., and Gruenberg, J. (1993). Cytoplasmic dynein-dependent vesicular transport from early to late endosomes. The Journal of cell biology 123, 1373–1387.

Boudewyn, L.C., and Walkley, S.U. (2018). Current concepts in the neuropathogenesis of mucolipidosis type IV. Journal of neurochemistry.

Bradford, M.M. (1976). A rapid and sensitive method for the quantitation of microgram quantities of protein utilizing the principle of protein-dye binding. Analytical biochemistry 72, 248–254.

Carstea, E.D., Morris, J.A., Coleman, K.G., Loftus, S.K., Zhang, D., Cummings, C., Gu, J., Rosenfeld, M.A., Pavan, W.J., Krizman, D.B., et al. (1997). Niemann-Pick C1 disease gene: homology to mediators of cholesterol homeostasis. Science 277, 228–231.

Chamoun, Z., Vacca, F., Parton, R.G., and Gruenberg, J. (2013). PNPLA3/adiponutrin functions in lipid droplet formation. Biol Cell 105, 219–233.

Chevallier, J., Chamoun, Z., Jiang, G., Prestwich, G.D., Sakai, N., Matile, S., Parton, R.G., and Gruenberg, J. (2008). Lysobisphosphatidic acid controls endosomal cholesterol levels. J Biol Chem 283, 27871–27880.

Dong, X.P., Wang, X., Shen, D., Chen, S., Liu, M., Wang, Y., Mills, E., Cheng, X., Delling, M., and Xu, H. (2009). Activating mutations of the TRPML1 channel revealed by proline-scanning mutagenesis. J Biol Chem 284, 32040–32052.

Encarnacao, M., Espada, L., Escrevente, C., Mateus, D., Ramalho, J., Michelet, X., Santarino, I., Hsu, V.W., Brenner, M.B., Barral, D.C., et al. (2016). A Rab3a-dependent complex essential for lysosome positioning and plasma membrane repair. The Journal of cell biology 213, 631–640.

Folch, J., Lees, M., and Sloane Stanley, G.H. (1957). A simple method for the isolation and purification of total lipides from animal tissues. J Biol Chem 226, 497–509.

Fukuda, M. (2013). Rab27 effectors, pleiotropic regulators in secretory pathways. Traffic (Copenhagen, Denmark) 14, 949–963.

Gong, X., Qian, H., Zhou, X., Wu, J., Wan, T., Cao, P., Huang, W., Zhao, X., Wang, X., Wang, P., et al. (2016). Structural Insights into the Niemann-Pick C1 (NPC1)-Mediated Cholesterol Transfer and Ebola Infection. Cell 165, 1467–1478.

Hartler, J., Trotzmuller, M., Chitraju, C., Spener, F., Kofeler, H.C., and Thallinger, G.G. (2011). Lipid Data Analyzer: unattended identification and quantitation of lipids in LC-MS data. Bioinformatics (Oxford, England) 27, 572–577.

Hissa, B., Pontes, B., Roma, P.M., Alves, A.P., Rocha, C.D., Valverde, T.M., Aguiar, P.H., Almeida, F.P., Guimaraes, A.J., Guatimosim, C., et al. (2013). Membrane cholesterol removal changes mechanical properties of cells and induces secretion of a specific pool of lysosomes. PLOS one 8, e82988.

Infante, R.E., and Radhakrishnan, A. (2017). Continuous transport of a small fraction of plasma membrane cholesterol to endoplasmic reticulum regulates total cellular cholesterol. eLife 6.

Infante, R.E., Radhakrishnan, A., Abi-Mosleh, L., Kinch, L.N., Wang, M.L., Grishin, N.V., Goldstein, J.L., and Brown, M.S. (2008). Purified NPC1 protein: II. Localization of sterol binding to a 240-amino acid soluble luminal loop. J Biol Chem 283, 1064–1075.

Kanerva, K., Uronen, R.L., Blom, T., Li, S., Bittman, R., Lappalainen, P., Peranen, J., Raposo, G., and Ikonen, E. (2013). LDL cholesterol recycles to the plasma membrane via a Rab8a-Myosin5b-actin-dependent membrane transport route. Dev Cell 27, 249–262.

Kobayashi, T., Beuchat, M.H., Chevallier, J., Makino, A., Mayran, N., Escola, J.M., Lebrand, C., Cosson, P., and Gruenberg, J. (2002). Separation and characterization of late endosomal membrane domains. J Biol Chem 277, 32157–32164.

Kobayashi, T., Beuchat, M.H., Lindsay, M., Frias, S., Palmiter, R.D., Sakuraba, H., Parton, R.G., and Gruenberg, J. (1999). Late endosomal membranes rich in lysobisphosphatidic acid regulate cholesterol transport. Nat Cell Biol 1, 113–118.

Kobayashi, T., Stang, E., Fang, K.S., de Moerloose, P., Parton, R.G., and Gruenberg, J. (1998). A lipid associated with the antiphospholipid syndrome regulates endosome structure and function. Nature 392, 193–197.

Kornfeld, S., and Mellman, I. (1989). The biogenesis of lysosomes. Annu Rev Cell Biol 5, 483–525.

Kristiana, I., Yang, H., and Brown, A.J. (2008). Different kinetics of cholesterol delivery to components of the cholesterol homeostatic machinery: implications for cholesterol trafficking to the endoplasmic reticulum. Biochim Biophys Acta 1781, 724–730.

Kwon, H.J., Abi-Mosleh, L., Wang, M.L., Deisenhofer, J., Goldstein, J.L., Brown, M.S., and Infante, R.E. (2009). Structure of N-terminal domain of NPC1 reveals distinct subdomains for binding and transfer of cholesterol. Cell 137, 1213–1224.

Laemmli, U.K. (1970). Cleavage of structural proteins during the assembly of the head of bacteriophage T4. Nature 227, 680–685.

Lange, Y., Ye, J., and Steck, T.L. (2004). How cholesterol homeostasis is regulated by plasma membrane cholesterol in excess of phospholipids. Proc Natl Acad Sci U S A 101, 11664–11667.

LaPlante, J.M., Sun, M., Falardeau, J., Dai, D., Brown, E.M., Slaugenhaupt, S.A., and Vassilev, P.M. (2006). Lysosomal exocytosis is impaired in mucolipidosis type IV. Mol Genet Metab 89, 339–348.

Laulagnier, K., Schieber, N.L., Maritzen, T., Haucke, V., Parton, R.G., and Gruenberg, J. (2011). Role of AP1 and Gadkin in the traffic of secretory endo-lysosomes. Mol Biol Cell 12, 2068–2082.

Li, X., Saha, P., Li, J., Blobel, G., and Pfeffer, S.R. (2016). Clues to the mechanism of cholesterol transfer from the structure of NPC1 middle lumenal domain bound to NPC2. Proc Natl Acad Sci U S A 113, 10079–10084.

Liscum, L. (2000). Niemann-Pick type C mutations cause lipid traffic jam. Traffic (Copenhagen, Denmark) 1, 218–225.

Liscum, L., and Faust, J.R. (1987). Low density lipoprotein (LDL)-mediated suppression of cholesterol synthesis and LDL uptake is defective in Niemann-Pick type C fibroblasts. J Biol Chem 262, 17002–17008.

Liu, B., Ramirez, C.M., Miller, A.M., Repa, J.J., Turley, S.D., and Dietschy, J.M. (2010). Cyclodextrin overcomes the transport defect in nearly every organ of NPC1 mice leading to excretion of sequestered cholesterol as bile acid. J Lipid Res 51, 933–944.

Liu, B., Turley, S.D., Burns, D.K., Miller, A.M., Repa, J.J., and Dietschy, J.M. (2009). Reversal of defective lysosomal transport in NPC disease ameliorates liver dysfunction and neurodegeneration in the npc1-/-mouse. Proc Natl Acad Sci U S A 106, 2377–2382.

Loizides-Mangold, U., David, F.P., Nesatyy, V.J., Kinoshita, T., and Riezman, H. (2012). Glycosylphosphatidylinositol anchors regulate glycosphingolipid levels. J Lipid Res 53, 1522–1534.

Luzio, J.P., Hackmann, Y., Dieckmann, N.M., and Griffiths, G.M. (2014). The biogenesis of lysosomes and lysosome-related organelles. Cold Spring Harbor perspectives in biology 6, a016840.

Marks, M.S., Heijnen, H.F., and Raposo, G. (2013). Lysosome-related organelles: unusual compartments become mainstream. Current opinion in cell biology 25, 495–505.

Martina, J.A., Diab, H.I., Lishu, L., Jeong, A.L., Patange, S., Raben, N., and Puertollano, R. (2014). The nutrient-responsive transcription factor TFE3 promotes autophagy, lysosomal biogenesis, and clearance of cellular debris. Sci Signal 7, ra9.

Medina, D.L., Fraldi, A., Bouche, V., Annunziata, F., Mansueto, G., Spampanato, C., Puri, C., Pignata, A., Martina, J.A., Sardiello, M., et al. (2011). Transcriptional activation of lysosomal exocytosis promotes cellular clearance. Dev Cell 21, 421–430.

Morel, E., and Gruenberg, J. (2007). The p11/S100A10 light chain of annexin A2 is dispensable for annexin A2 association to endosomes and functions in endosomal transport. PLOS one 2, e1118.

Naureckiene, S., Sleat, D.E., Lackland, H., Fensom, A., Vanier, M.T., Wattiaux, R., Jadot, M., and Lobel, P. (2000). Identification of HE1 as the second gene of Niemann-Pick C disease. Science 290, 2298–2301.

Ory, D.S., Ottinger, E.A., Farhat, N.Y., King, K.A., Jiang, X., Weissfeld, L., Berry-Kravis, E., Davidson, C.D., Bianconi, S., Keener, L.A., et al. (2017). Intrathecal 2-hydroxypropyl-beta-cyclodextrin decreases neurological disease progression in Niemann-Pick disease, type C1: a nonrandomised, open-label, phase 1-2 trial. Lancet 390, 1758–1768.

Ostrowski, M., Carmo, N.B., Krumeich, S., Fanget, I., Raposo, G., Savina, A., Moita, C.F., Schauer, K., Hume, A.N., Freitas, R.P., et al. (2010). Rab27a and Rab27b control different steps of the exosome secretion pathway. Nat Cell Biol 12, 19-30; sup pp 11–13.

Pankiv, S., Alemu, E.A., Brech, A., Bruun, J.A., Lamark, T., Overvatn, A., Bjorkoy, G., and Johansen, T. (2010). FYCO1 is a Rab7 effector that binds to LC3 and PI3P to mediate microtubule plus end-directed vesicle transport. The Journal of cell biology 188, 253–269.

Patterson, M.C., Mengel, E., Vanier, M.T., Schwierin, B., Muller, A., Cornelisse, P., and Pineda, M. (2015). Stable or improved neurological manifestations during miglustat therapy in patients from the international disease registry for Niemann-Pick disease type C: an observational cohort study. Orphanet journal of rare diseases 10, 65.

Pentchev, P.G., Comly, M.E., Kruth, H.S., Vanier, M.T., Wenger, D.A., Patel, S., and Brady, R.O. (1985). A defect in cholesterol esterification in Niemann-Pick disease (type C) patients. Proc Natl Acad Sci U S A 82, 8247–8251.

Pfisterer, S.G., Peranen, J., and Ikonen, E. (2016). LDL-cholesterol transport to the endoplasmic reticulum: current concepts. Current opinion in lipidology 27, 282–287.

Pu, J., Schindler, C., Jia, R., Jarnik, M., Backlund, P., and Bonifacino, J.S. (2015). BORC, a multisubunit complex that regulates lysosome positioning. Dev Cell 33, 176–188.

Raiborg, C., Wenzel, E.M., Pedersen, N.M., Olsvik, H., Schink, K.O., Schultz, S.W., Vietri, M., Nisi, V., Bucci, C., Brech, A., et al. (2015). Repeated ER-endosome contacts promote endosome translocation and neurite outgrowth. Nature 520, 234–238.

Ran, F.A., Hsu, P.D., Wright, J., Agarwala, V., Scott, D.A., and Zhang, F. (2013). Genome engineering using the CRISPR-Cas9 system. Nature protocols 8, 2281–2308.

Reddy, A., Caler, E.V., and Andrews, N.W. (2001). Plasma membrane repair is mediated by Ca(2+)-regulated exocytosis of lysosomes. Cell 106, 157–169.

Rocha, N., Kuijl, C., van der Kant, R., Janssen, L., Houben, D., Janssen, H., Zwart, W., and Neefjes, J. (2009). Cholesterol sensor ORP1L contacts the ER protein VAP to control Rab7-RILP-p150 Glued and late endosome positioning. The Journal of cell biology 185, 1209–1225.

Rosa-Ferreira, C., and Munro, S. (2011). Arl8 and SKIP act together to link lysosomes to kinesin-1. Dev Cell 21, 1171–1178.

Rosenbaum, A.I., and Maxfield, F.R. (2011). Niemann-Pick type C disease: molecular mechanisms and potential therapeutic approaches. Journal of neurochemistry 116, 789–795.

Rosenbaum, A.I., Zhang, G., Warren, J.D., and Maxfield, F.R. (2010). Endocytosis of beta-cyclodextrins is responsible for cholesterol reduction in Niemann-Pick type C mutant cells. Proceedings of the National Academy of Sciences of the United States of America 107, 5477–5482.

Samie, M.A., and Xu, H. (2014). Lysosomal exocytosis and lipid storage disorders. J Lipid Res 55, 995–1009.

Saridaki, T., Nippold, M., Dinter, E., Roos, A., Diederichs, L., Fensky, L., Schulz, J.B., and Falkenburger, B.H. (2018). FYCO1 mediates clearance of alpha-synuclein aggregates through a Rab7-dependent mechanism. Journal of neurochemistry.

Savina, A., Vidal, M., and Colombo, M.I. (2002). The exosome pathway in K562 cells is regulated by Rab11. J Cell Sci 115, 2505–2515.

Scott, C.C., Vossio, S., Vacca, F., Snijder, B., Larios, J., Schaad, O., Guex, N., Kuznetsov, D., Martin, O., Chambon, M., et al. (2015). Wnt directs the endosomal flux of LDL-derived cholesterol and lipid droplet homeostasis. EMBO Rep 16, 741–752.

Sheriff, S., Du, H., and Grabowski, G.A. (1995). Characterization of lysosomal acid lipase by site-directed mutagenesis and heterologous expression. J Biol Chem 270, 27766–27772.

Simons, K., and Gruenberg, J. (2000). Jamming the endosomal system: lipid rafts and lysosomal storage diseases. Trends in cell biology 10, 459–462.

Sobo, K., Chevallier, J., Parton, R.G., Gruenberg, J., and van der Goot, F.G. (2007). Diversity of raftlike domains in late endosomes. PLOS one 2, e391.

Vance, J.E., and Peake, K.B. (2011). Function of the Niemann-Pick type C proteins and their bypass by cyclodextrin. Current opinion in lipidology 22, 204–209.

Vanier, M.T. (2010). Niemann-Pick disease type C. Orphanet journal of rare diseases 5, 16.

Vite, C.H., Bagel, J.H., Swain, G.P., Prociuk, M., Sikora, T.U., Stein, V.M., O’Donnell, P., Ruane, T., Ward, S., Crooks, A., et al. (2015). Intracisternal cyclodextrin prevents cerebellar dysfunction and Purkinje cell death in feline Niemann-Pick type C1 disease. Sci Transl Med 7, 276ra226.

Wang, M.L., Motamed, M., Infante, R.E., Abi-Mosleh, L., Kwon, H.J., Brown, M.S., and Goldstein, J.L. (2010). Identification of surface residues on Niemann-Pick C2 essential for hydrophobic handoff of cholesterol to NPC1 in lysosomes. Cell Metab 12, 166–173.

Xu, M., Liu, K., Swaroop, M., Porter, F.D., Sidhu, R., Firnkes, S., Ory, D.S., Marugan, J.J., Xiao, J., Southall, N., et al. (2012). delta-Tocopherol reduces lipid accumulation in Niemann-Pick type C1 and Wolman cholesterol storage disorders. J Biol Chem 287, 39349–39360.

Xu, S., Benoff, B., Liou, H.L., Lobel, P., and Stock, A.M. (2007). Structural basis of sterol binding by NPC2, a lysosomal protein deficient in Niemann-Pick type C2 disease. J Biol Chem 282, 23525–23531.

Zhong, X.Z., Sun, X., Cao, Q., Dong, G., Schiffmann, R., and Dong, X.P. (2016). BK channel agonist represents a potential therapeutic approach for lysosomal storage diseases. Sci Rep 6, 33684.

